# Distinct roles of cortical layer 5 subtypes in associative learning

**DOI:** 10.1101/2025.01.07.631500

**Authors:** Sara Moberg, Michele Garibbo, Camille Mazo, Ariel Gilad, Dietmar Schmitz, Rui Ponte Costa, Matthew E. Larkum, Naoya Takahashi

## Abstract

Adaptive behavior is critically dependent on associative learning, where environmental cues are linked with subsequent positive or negative outcomes. In mammals, primary neocortical sensory areas serve as pivotal nodes in this process, processing stimuli and distributing information to cortical and subcortical networks. Layer 5 (L5) of the cortex comprises two types of pyramidal projection neurons—intratelencephalic (IT) and extratelencephalic (ET) neurons—each with distinct downstream targets. Despite the crucial function of L5 as a main output node of the cortex, the specific contributions of these L5 neuronal subtypes to associative learning remain poorly understood. In the present study, by leveraging transgenic mouse lines, we distinguished IT and ET neurons in the primary somatosensory cortex and examined their roles in a whisker-based frequency-discrimination learning task. Longitudinal two-photon calcium imaging revealed distinct response characteristics between IT and ET neurons throughout learning. Interestingly, the activity of IT neurons hardly changed over the five days of learning, while the activity of ET neurons developed robustly. Furthermore, IT neurons appeared to show stimuli encoding from the beginning, whereas the ET neurons became increasingly responsive to stimuli associated with reward. Chemogenetic silencing of either IT or ET neurons both impaired learning, but in strikingly distinct ways, each associated with a different phase of learning. By modeling the response characteristics of IT and ET neurons using a reinforcement learning framework, we show that IT neurons primarily encode sensory stimuli, and their representations are critical for forming stimulus-reward associations. ET neurons instead represent the value of the stimulus, used for refining behavior. Thus, our results delineate the distinct roles of L5 IT and ET neurons, underscoring their integral and complementary contributions to associative learning.

## INTRODUCTION

Associative learning is essential for selecting appropriate behaviors by linking stimuli to different outcomes. The learning process consists of multiple components, including acquisition and refinement, and involves diverse neural circuits across the brain that function in concert^1,2^. Primary sensory cortical areas are crucial for first processing and analyzing sensory stimuli, but then also for linking these stimuli to behaviorally relevant outcomes, such as rewards or threats^3–6^. During stimulus-reward associative learning, neurons in these areas alter their response patterns to stimuli based on their relevance to rewards^3,7–9^. These learning-associated changes in neuronal activity are thought to then be transmitted to other brain regions outside the sensory cortices. While recent studies have begun to elucidate the contribution to learning of cortical neurons projecting to distant brain regions^5^, little is known about the activity changes these projection neurons undergo during learning and their functional roles throughout the learning process.

Layer 5 (L5) serves as the main output layer of sensory cortices and consists of two major groups of outward projecting pyramidal neurons: intratelencephalic (IT) and extratelencephalic (ET) neurons. IT neurons project bilaterally to other cortical areas and the striatum, while ET neurons project predominantly to subcortical areas in the ipsilateral hemisphere, including the striatum, higher-order thalamic nuclei, midbrain and pons^10–12^. Recent advances in transgenic^13^ and retrograde-viral targeting^14^ technology have begun to shed light on the distinct contribution of each projection neuronal subtype to cortical function and information processing, such as sensory detection and discrimination^4,5,15,16^, decision making^17^, and movement control^18–21^. But as yet, the question still remains as to how IT and ET neurons contribute to learning.

The primary somatosensory cortex (S1) has been shown to engage in vibrotactile discrimination in humans^22^, primates^23^, and rodents^24^. In the present study, we sought to determine how the activity patterns of S1 L5 projection neuronal subtypes evolve during learning of vibrotactile discrimination, specifically in the context of stimulus-reward associations. To study the learning process, we designed an appetitive Pavlovian conditioning task, where head-restrained mice were trained to distinguish between two vibratory stimuli delivered to their whiskers with different frequencies, with one stimulus paired with a water reward and the other not. Utilizing transgenic mouse lines, we selectively targeted and tracked the activity of either L5 IT or ET neuronal populations in S1 during the learning process. Using *in vivo* two-photon imaging, we measured calcium activity in the apical dendritic trunks, which highly correlates with somatic activity of L5 pyramidal neurons^25,26^, and found distinct response patterns between IT and ET neurons to stimuli and rewards, along with changes in these patterns during learning. By chemogenetically silencing selective L5 subtypes, we found that both IT and ET neurons are critical for learning, with each subtype contributing to distinct aspects of the learning process. To further elucidate the specific roles of IT and ET neurons, we developed a theoretical model inspired by classical reinforcement learning frameworks and based on their distinctive response properties. The model suggests that IT neurons encode an ‘unsupervised’ model of the sensory world, which can then be used to initiate the formation of associations between stimuli and rewards. In contrast, the model suggests that ET neurons are pivotal in the subsequent refinement of learned responses, thereby enhancing the discriminability between rewarding and non-rewarding stimuli.

## RESULTS

### Whisker stimulus-reward associative learning depends on S1

To study reward-based associative learning in mice, we developed a Pavlovian trace conditioning task in which head-restrained mice were trained to associate one of two stimuli, i.e. a conditioned stimulus (CS+) vs. a non-conditioned stimulus (CS-), with water reward (**Fig. 1a**). The CS+ consisted of C2-whisker stimulation at 10 Hz for 1s, followed by a short delay (trace interval; 0.5 s) before the delivery of a reward. For CS-trials, the same whisker was stimulated at 5 Hz with no reward. Over multiple daily sessions of training, the mice gradually learned to discriminate between the two stimuli, as shown by a selective increase in the frequency of anticipatory licks to CS+ (i.e., licks from stimulus onset to the time of reward delivery) (**Fig. 1b**). The task performance, i.e., CS discriminability, was quantified as the area under a receiver operating characteristic curve (auROC) calculated on anticipatory licks in CS+ vs. CS-trials for each session (**Fig. 1c**)^27,28^. Within five days, 18 out of 20 mice reached a proficient level (auROC > 0.7) to discriminate the stimuli (**Fig. 1d**). Learning was not contingent on the specific frequencies of conditioned stimuli used as mice could successfully learn the task where 5 Hz stimulation was set as CS+ and 10 Hz stimulation as CS-(**Fig. S1**).

**Figure 1.**
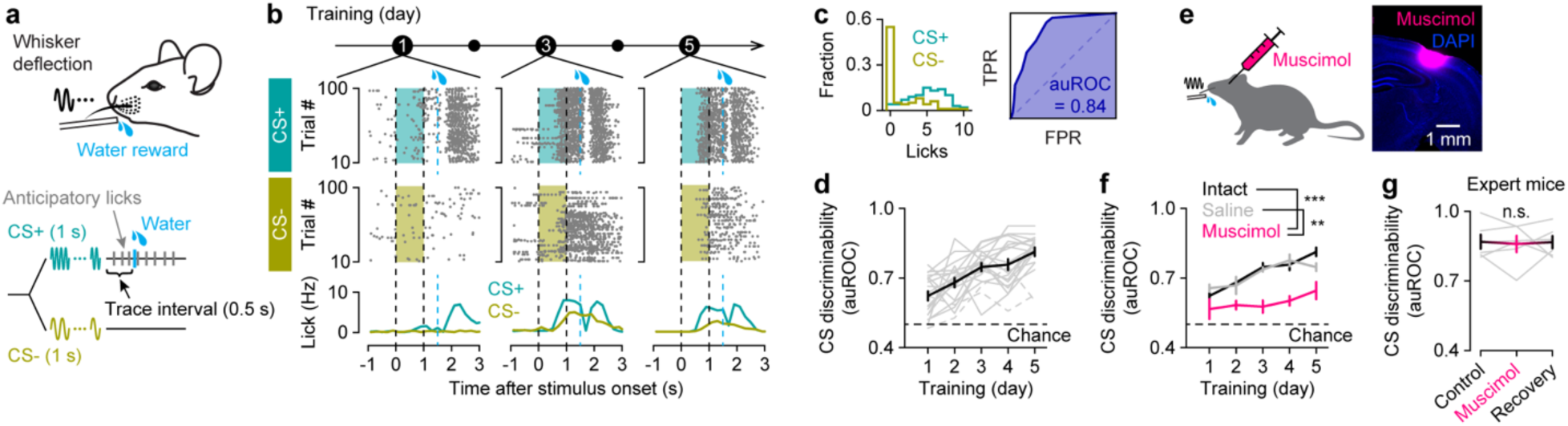
Whisker stimulus-reward associative learning depends on S1. (a) Behavioral task design. Mice were exposed to two different whisker stimuli of different frequencies: one stimulus (CS+, 1 s) followed by a reward after a short delay (trace interval, 0.5 s), and the other (CS-, 1 s) left unrewarded. (b) Example training sessions (Days 1, 3 and 5) from a mouse showing anticipatory and consummatory licking during CS+ and CS-trials. Licking activities are shown in raster plots (top) and histograms (bottom). Top, raster plot with trial sorted according to stimulus type (CS+ vs. CS-). Shaded areas, stimulus presentation; black dashed lines, stimulus onset and offset; blue dashed lines, reward onset for CS+ trials. (c) Left, distribution of the anticipatory lick counts during CS+ (blue) and CS-trials (yellow) of an example session (Day 5 in **b**). Right, the area under the receiver operating characteristic (auROC), scoring behavioral performance. TPR: true positive rate; FPR: false positive rate. (d) Evolution of CS discrimination performance over five days of training (*n* = 20 mice; *p* = 8.1 × 10^-12^, *F* = 21.32; one-way repeated-measure ANOVA). Gray lines, individual mice; dashed gray lines, mice that did not learn the task; black dashed line, chance level of behavioral discrimination between CS. Data are presented as mean ± SEM. (e) Coronal section of S1, showing the diffusion of muscimol injected in the C2 barrel column. (f) Learning trajectory of mice with intact S1 (*n* = 20 mice), mice with saline injection (*n* = 6 mice), and mice with muscimol injection (*n* = 6 mice; *p* = 1.2 × 10^-4^, *F* = 12.48; two-way repeated-measure ANOVA with post hoc Tukey-Kramer test). \*\**p* < 0.01, \*\*\**p* < 0.001. Data are presented as mean ± SEM. (g) Behavioral performance of expert mice injected with muscimol (*n* = 6 mice; *p* = 0.95, *F* = 0.055; one-way repeated-measure ANOVA). Data are presented as mean ± SEM; gray lines, individual mice.

A recent report suggests that learning a simple tactile detection task may not require S1^29^. To test whether our learning paradigm depends on S1, we pharmacologically inactivated it during training. Prior to each training session, muscimol, a GABA_A_-receptor agonist, was locally injected into the C2 barrel column of S1, contralaterally to the stimulated C2 whisker (**Fig. 1e**). Silencing S1 disrupted learning compared to mice with intact S1 or S1 injected with saline (**Fig. 1f**). In contrast, silencing S1 in expert mice did not affect the behavioral performance (**Fig. 1g**). Collectively, these results demonstrate that S1 activity facilitates learning of the task, but once learned, S1 is not necessary for the expression of conditioned responses.

### IT neurons respond primarily to whisker stimuli while ET neurons are sensitive to reward-related signals

To investigate the responses of S1 neurons to stimuli and rewards, and how they are modulated during learning, we used *in vivo* two-photon calcium imaging to monitor neuronal activity over time before, during and after learning. We focused on the two main long-range projection neurons in L5 of S1, IT and ET neurons. Two Cre-driver transgenic mouse lines, Tlx3-Cre and Sim1-Cre, were used to distinguish between IT and ET neurons, respectively (**Fig. 2a**)^13^. A Cre-dependent viral expression strategy yielded selective labeling of L5 IT neurons in Tlx3-Cre mice and ET neurons in Sim1-Cre mice, as previously reported (**Fig. 2b**)^13,16,19,20^.

**Figure 2.**
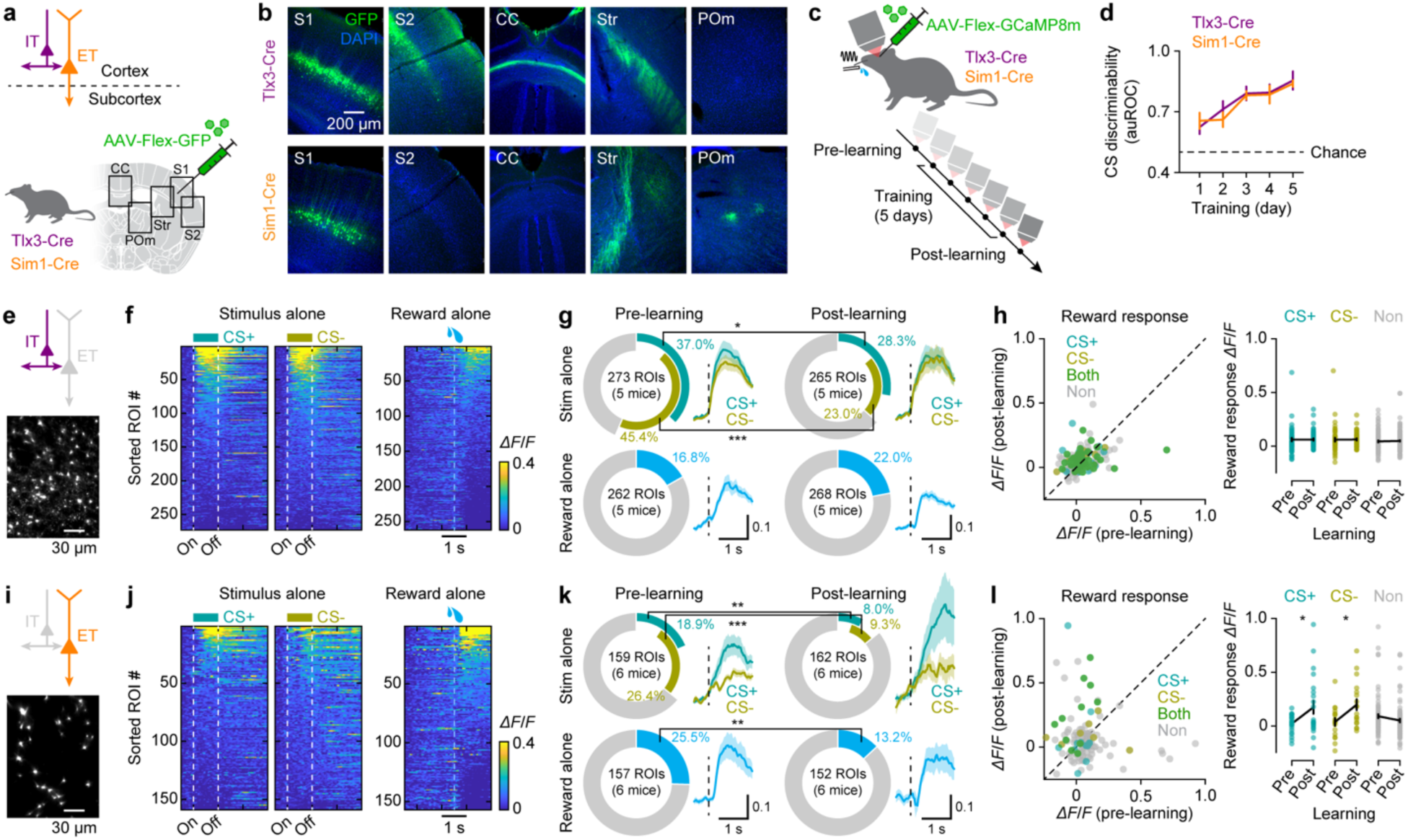
Distinct response patterns of L5 neuronal subtypes to sensory stimuli and reward. (a) Schematic showing the genetic and viral intersectional strategy, targeting L5 IT neurons in Tlx3-Cre mice and ET neurons in Sim1-Cre mice. (b) Images from coronal sections from S1 of a Tlx3-Cre mouse (top) and Sim1-Cre mouse (bottom) injected with AAV-Flex-GFP, showing the injection site in S1 and labeled axons in the secondary somatosensory cortex (S2), corpus callosum (CC), striatum (Str), and posterior medial thalamic nucleus (POm). (c) Longitudinal two-photon calcium imaging from Tlx3-Cre and Sim1-Cre mice injected with AAV-Flex-jGCaMP8m in S1 before, during, and after five days of training. (d) Behavioral performance of Tlx3-Cre and Sim1-Cre mice used in imaging experiments (*n* = 5 Tlx3-Cre mice and 6 Sim1-Cre mice; *p* = 0.96, *F* = 0.0032; two-way repeated measure ANOVA). Data are presented as mean ± SEM. (e) An example field of view of IT neuronal dendrites. (f) Heatmaps of IT neuronal responses to stimulus alone (left, CS+ responses; middle, CS-responses; *n* = 273 neurons) and reward alone (right, *n* = 262 neurons) before learning (*n* = 5 mice). ROIs in each heatmap are sorted by their mean response amplitudes within 1.5 s of stimulus or reward onset. (g) Top, fraction of IT neurons responding to CS+ or CS-before and after learning (\**p* = 0.032 for CS+; \*\*\**p* = 4.5 × 10^-8^ for CS-; 2 × 2 χ^2^ test). Inset, average responses of CS-responding IT neurons (mean ± SEM). Dashed line, stimulus onset. Bottom, fraction of IT neurons responding to reward before and after learning (*p* = 0.13; 2 × 2 χ^2^ test). (h) Left, scatter plot showing IT neurons’ responses to reward before and after learning. Each circle represents an individual neuron. Colors represent neurons’ CS selectivity before learning (blue, CS+ only; yellow, CS-only, green, both stimuli). Right, IT neurons’ response amplitudes to reward before vs. after learning (mean ± SEM), separated by their CS selectivity before learning (*n* = 58 neurons responding to CS+, *p* = 0.95; *n* = 51 neurons responding to CS-, *p* = 0.91; *n* = 104 non-responding neurons, *p* = 0.24; two-sample *t*-test). (**i–l**) Same as **e–h** but for ET neurons. (**j**) *n* = 159 neurons for stimulus alone, *n* = 157 neurons for reward (*n* = 6 mice). (**k**) Top, \*\**p* = 0.0043 for CS+, \*\*\**p* = 5.8 × 10^-5^ for CS-. Bottom, \*\**p* = 0.0062. (**l**) *n* = 22 neurons responding to CS+, \**p* = 0.020; *n* = 18 neurons responding to CS-, \**p* = 0.013; *n* = 96 non-responding neurons, *p* = 0.11.

Using these mouse lines, we expressed a genetically-encoded calcium indicator, jGCaMP8m^30^, in IT and ET neurons. To facilitate stable longitudinal recordings, we imaged apical dendritic trunks as a proxy for global activity as activities in apical trunks and somata have been shown to be highly correlated^25,26^. The same field of view (FOV) in the C2 barrel column was imaged for seven consecutive days; five days during the course of learning as well as one imaging session before and one imaging session after learning to record the neurons’ response profiles to stimulus alone and reward alone (**Fig. 2c**). Tlx3-Cre and Sim1-Cre transgenic mice learned the task equally well over the course of five days (**Fig. 2d**). On average, the FOV (178.3 × 178.3 µm^2^) contained 56 IT neurons (34–83 neurons, *n* = 5 mice; **Fig. 2e**) or 28 ET neurons (11–45 neurons, *n* = 6 mice; **Fig. 2i**).

When comparing the responses of IT and ET neurons in passive mice, where stimuli and rewards were presented separately in different trial blocks (**Fig. 2f, j**), we observed striking differences between the two subtypes. A larger fraction of IT neurons responded to whisker stimuli compared to ET neurons (pre-learning: 56.41% vs. 35.85%, *n* = 273 IT neurons vs. 159 ET neurons; *p* = 3.7 × 10^-5^, χ^2^ test = 17.00; **Fig. 2g, k**). Subpopulations of IT neurons exhibited CS selectivity; however, a notable fraction of neurons responded to both CS+ and CS-(**Fig. 2g**). Regardless of subtype, responsiveness to stimuli tended to decrease after learning.

In contrast, a greater proportion of ET neurons responded to rewards compared to IT neurons (pre-learning: 16.79% vs. 25.48%, *n* = 262 IT neurons vs.157 ET neurons; *p* = 0.032, χ^2^ test = 4.62; **Fig. 2g, k**). Similar to their stimulus responsiveness, ET neurons’ reward responsiveness also declined after learning. Interestingly, ET, but not IT, neurons that initially responded to stimuli showed increased responsiveness to rewards after learning (**Fig. 2h, l**). This suggests a learning-induced shift in the response properties of ET neurons, where stimulus-responding ET neurons preferentially develop increased sensitivity to rewards through the learning process. Collectively, our results show that IT neurons primarily responded to stimuli, while ET neurons were more responsive to rewards.

### Learning reshapes ET neuronal responses, with little change in IT neurons

To assess how IT and ET neurons change their activity patterns during learning, we recorded from the same mice over successive daily training sessions. The response patterns of IT neurons remained stable throughout the learning period (**Fig. 3a, b**). In contrast to outside the training (**Fig. 2**), where we observed a decrease in stimuli-responsive neurons, the fraction of stimulus-responding IT neurons did not change throughout learning (**Fig. 3c, left**), nor did their response amplitudes (**Fig. 3c, right**). Similarly to naive mice (**Fig. 2g**), the IT neurons responding to CS+ and CS-largely overlapped (green; **Fig. 3c, left**), with only a subset of neurons showing stimulus-specific responses (blue and yellow; **Fig. 3c, left**). ET neurons, on the other hand, showed a robust response to rewards on Day 1. Their response patterns gradually changed over the training period, parallel to the evolution of anticipatory lick patterns in response to the stimuli (**Fig. 3e, f**). Learning altered the proportion of engaged ET neurons as well as their response amplitudes (**Fig. 3g**). The response amplitudes for both stimuli significantly increased from the beginning to the end of training (**Fig. 3g, right**). Importantly, ET neuronal activity emerged with anticipatory licking but did not merely reflect lick-related motor activity (**Fig. S2**). Instead, it represented a response to reward-predicting stimuli. Together, these results suggest stable, sensory representations in IT neurons and learning-induced increase in ET neuron activity.

**Figure 3.**
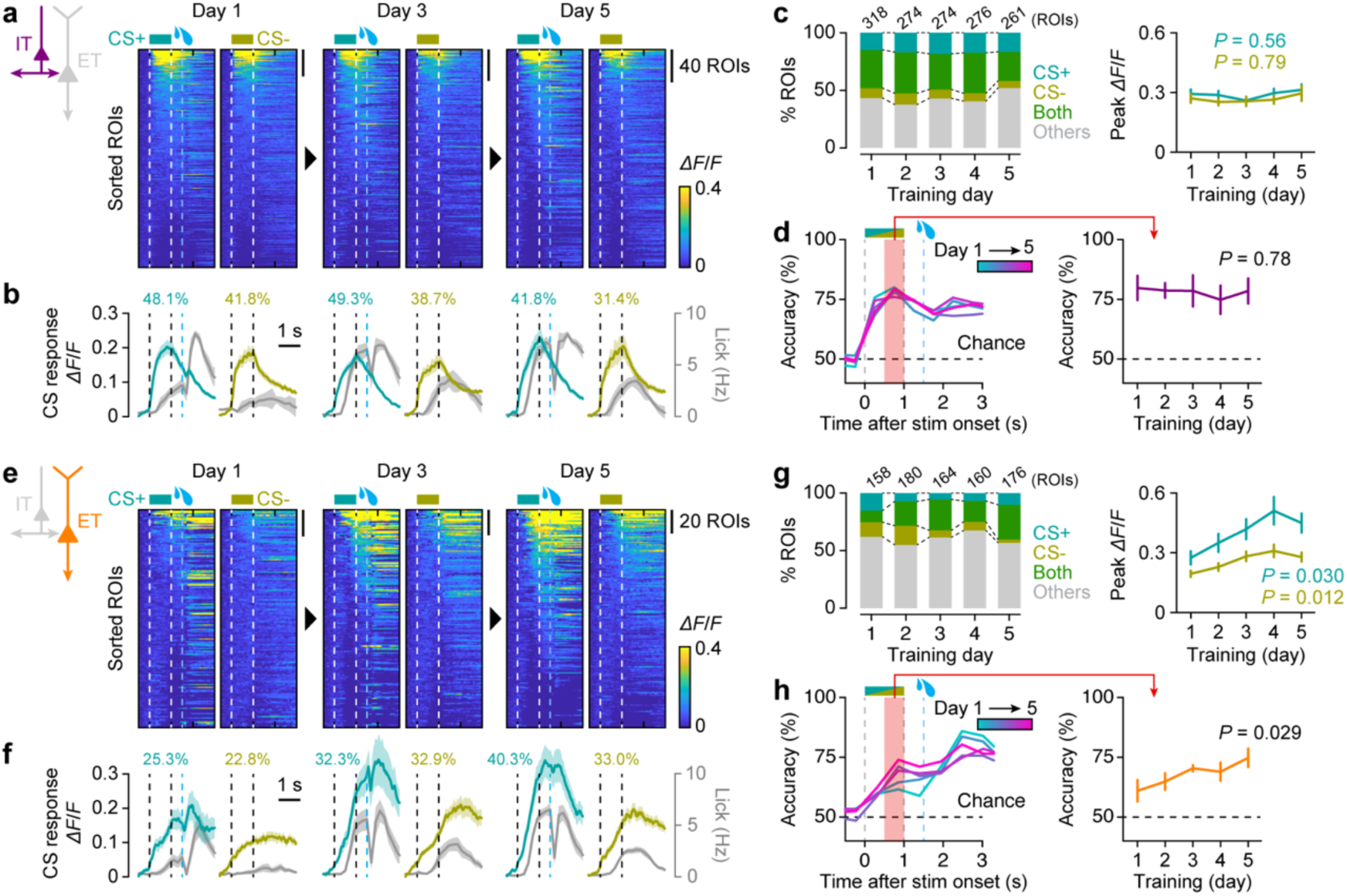
L5 IT neurons maintain robust stimulus encoding, while ET neurons develop reward-expectation responses during learning. (a) Heatmaps of trial-averaged responses of IT neurons during learning, sorted based on the mean response amplitudes from the stimulus onset until the reward onset at each day (*n* = 5 mice). (b) Colored trace, calcium responses average across IT neurons responding to CS+ (blue) or responding to CS-(yellow). Inset, the numbers indicating the fractions of IT neurons responding to CS+ or responding to CS-. Gray trace, average lick rates. Data are presented as mean ± SEM (c) Left, fractions of IT neurons responding to CS+ only (blue), CS-only (yellow), or both stimuli (green) (*p* = 0.14, χ^2^ test = 17.24; 4 × 5 χ^2^ test). Right, peak response across the 5 days to CS+ trials (blue; *F* = 0.75; one-way ANOVA) and CS-trials (yellow; *F* = 0.43; one-way ANOVA). Data are presented as mean ± SEM. (d) Left, SVM decoder performance in classifying the trial types (CS+ or CS-) at different time points within the task over training Days 1–5, based on IT neuronal responses (*n* = 5 mice). Right, decoder accuracies for the time window marked in pink on the left (*F* = 0.44; one-way repeated measure ANOVA). Data are presented as mean ± SEM. (**e–h**) Same as **a–d** but for ET neurons (*n* = 6 mice). (**g**) Left, fractions of ET neurons responding to CS+ only (blue), CS-only (yellow), or both stimuli (green) (*p* = 1.8 × 10^-7^, χ^2^ test = 55.02; 4 × 5 χ^2^ test). Right, peak response across the 5 days to CS+ trials (blue; *F* = 2.72; one-way ANOVA) and CS-trials (yellow; *F* = 3.29; one-way ANOVA). (**h**) Right, *F* = 3.37; one-way repeated measure ANOVA. Data are presented as mean ± SEM.

Since mice improved their performance in discriminating between CS+ and CS-through learning, we next sought to investigate how the activity of each neuronal subtype developed during learning. To investigate this, we performed a population-decoding analysis using a linear support vector machine (SVM) decoder to classify stimulus types on a trial-by-trial basis. Stimulus type was decoded throughout the trial using time-binned population activity (bin = 0.5 s) for Days 1–5. There was a consistent sharp rise in the decoding accuracy of IT neurons after the stimulus presentation and the accuracy remained high throughout the trial (**Fig. 3d**). These results reveal a robust populational representation of individual stimuli in IT neurons that change little during learning. In contrast, ET neurons showed high CS discriminability in the reward time window at the beginning of learning, while a second peak gradually appeared during the stimulus presentation time window over the training period (**Fig. 3h**). These results align with the observation that many ET neurons were responsive to reward before and at the early phase of learning (Day 1), and that an increase in activity associated with reward expectation emerged as learning progressed (**Fig. 3h**).

Given the distinct activity patterns of IT and ET neuronal populations, we wondered how individual neuronal activities evolve during learning. To assess this, we next analyzed neurons consistently identifiable across the five-day training period and tracked daily changes in their activity (**Fig. 4a, c**). Individual IT neurons exhibited stable response patterns throughout the learning period. A largely consistent neuronal population was activated by each stimulus from day to day (**Fig. 4a**). To further assess the response stability, we calculated the Pearson correlation of session-averaged activity traces for each neuron across different learning days and averaged these correlations across the population. This analysis confirmed that IT neuronal responses remained highly stable across days for both trial types (**Fig. 4b**). In contrast, ET neurons displayed more variable response patterns, with their activity changing from day to day (**Fig. 4c**), resulting in lower inter-day correlations (**Fig. 4d**). However, these correlations tended to gradually increase over time, suggesting that ET neuronal responses became more consistent. Together, these results suggest that individual IT neurons maintain stable activity that is unaffected by learning, while individual ET neurons undergo dynamic changes, with their activity patterns becoming more consistent as learning progresses.

**Figure 4.**
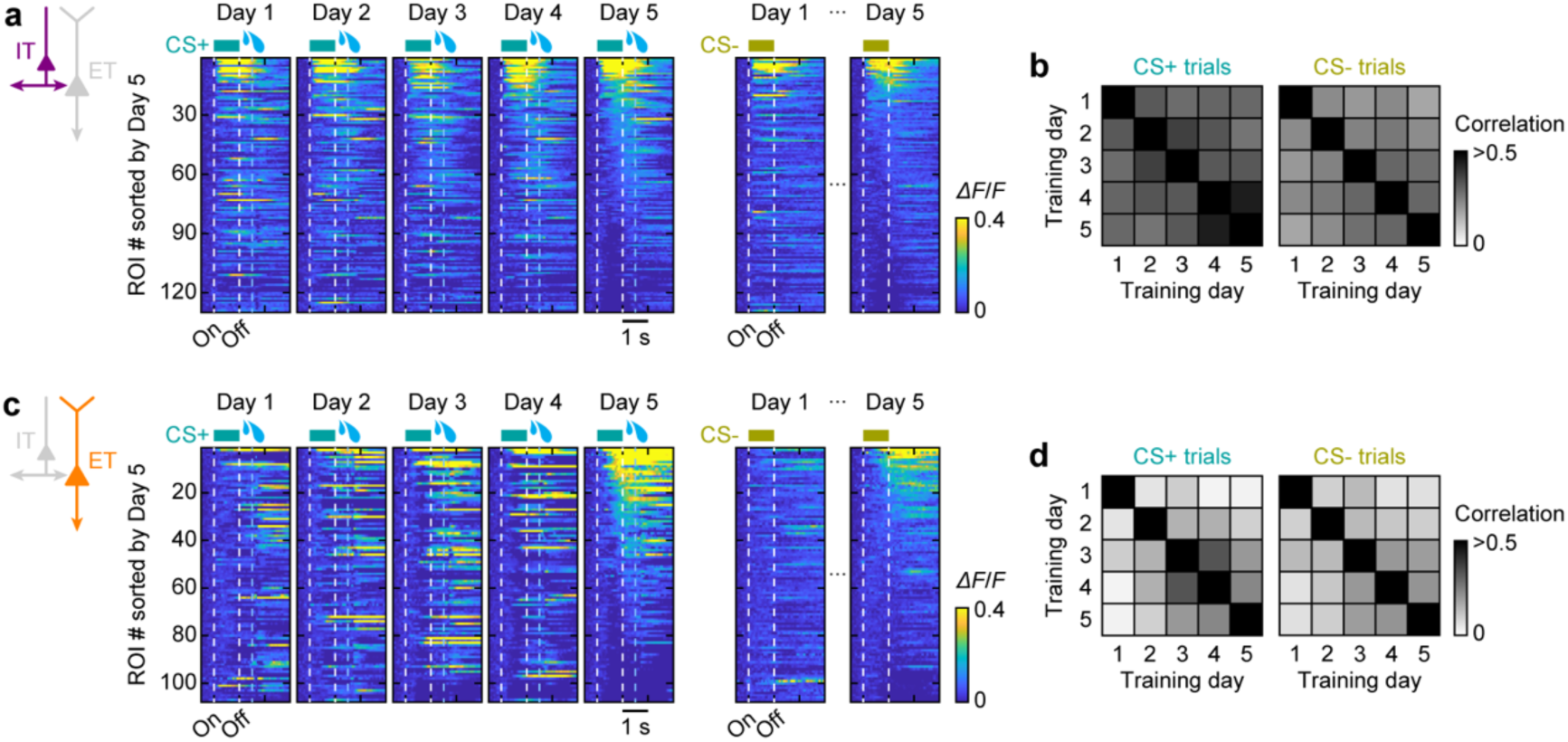
Static IT neuronal encoding and adaptive ET neuronal responses throughout learning. (a) Heatmaps of trial-averaged responses of IT neurons during learning, sorted based on the calcium response amplitudes at Day 5. (b) Pairwise Pearson’s correlation coefficient between training days of matched single neurons’ trial-averaged traces (*n* = 130 tracked IT neurons from *n* = 5 mice). The values were averaged across all neurons for each mouse and then averaged across mice. (**c, d**) Same as **a** and **b** but for ET neurons. (**d**) *n* = 108 tracked ET neurons from *n* = 6 mice.

### IT and ET neurons contribute to different aspects of learning

The differences in responsiveness between IT and ET neurons during learning raise the question of how these two neuronal subtypes uniquely contribute to behavioral learning. We therefore examined the specific functional roles of IT and ET neuronal populations during the associative learning task. We silenced each neuronal subtype in the C2 barrel by injecting a viral vector conditionally expressing an inhibitory designer receptor exclusively activated by designer drug (DREADD), hM4Di, into Tlx3-Cre or Sim1-Cre mice (**Fig. 5a–c**)^31^. Prior to each training session, a DREADD ligand, clozapine-N-oxide (CNO), was systemically injected into the mice, thus selectively inhibiting IT or ET neurons. Silencing either neuronal subtype led to a significant impairment in learning compared to control mice with intact S1 (**Fig. 5d**). The disruption of learning was not due to CNO injections alone as the learning remained intact when injecting CNO in mice without hM4Di expression (**Fig. S3**).

**Figure 5.**
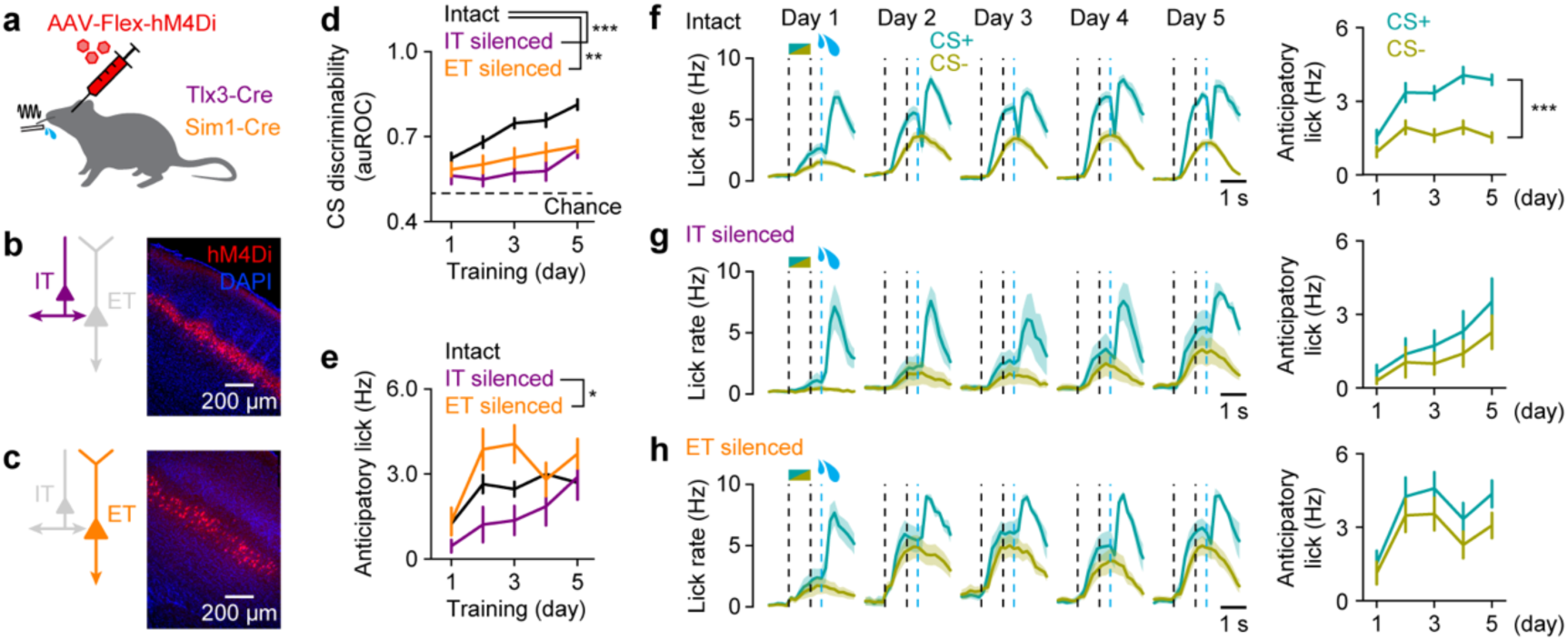
Silencing IT or ET neurons disrupts learning, but with different effects. (a) Targeted chemogenetic silencing of IT or ET neurons in S1 in Tlx3-Cre or Sim1-Cre mice, respectively. (b) Example image of a coronal section of S1 where IT neurons expressed hM4Di-mcherry. (c) Same as **b** but for ET neurons. (d) Behavioral performance of mice with S1 intact (*n* = 20 mice), silenced IT neurons (*n* = 6 mice), ET silenced ET neurons (*n* = 6 mice; *p* = 1.9 × 10^-5^, *F* = 16.15; two-way repeated-measure ANOVA with post hoc Tukey-Kramer test). Data are presented as mean ± SEM. \*\**p* < 0.01, \*\*\**p* < 0.001. (e) Mean anticipatory lick rates of mice with S1 intact (*n* = 20 mice), silenced IT neurons (*n* = 6 mice), and silenced ET neurons (*n* = 6 mice; *p* = 0.034, *F* = 3.80; two-way repeated-measure ANOVA with post hoc Tukey-Kramer test). Data are presented as mean ± SEM. \**p* < 0.05. (f) Left, lick rates for CS+ (blue) and CS-(yellow) average across mice for Days 1–5 (*n* = 20 mice). Right, evolution of mean anticipatory lick rates during learning (*p* = 1.5 × 10^-4^, *F* = 6.06; two-way repeated-measure ANOVA). Dashed lines represent CS onset and offset and reward onset. Data are presented as mean ± SEM. \*\*\**p* < 0.001. (g) Same as **f** but for mice with silenced IT neurons (*p* = 0.77, *F* = 0.45; two-way repeated-measure ANOVA). (h) Same as **f** but for mice with silenced ET neurons (*p* = 0.82, *F* = 0.38; two-way repeated-measure ANOVA).

Interestingly, the progression of overall anticipatory lick rates significantly differed between IT and ET neuron silencing (**Fig. 5e**). Under closer observation, during learning in control mice, anticipatory lick rates for CS+ increased as learning progressed, while lick rates for CS-increased during the first two days and then trended to decline (**Fig. 5f**). Inactivating IT neurons caused a general reduction in anticipatory lick rates for both CS+ and CS-, with a more pronounced decrease for CS+ (**Fig. S4a**), ultimately eliminating the difference between the two (**Fig. 5g**). Conversely, ET neuron inactivation resulted in a sharp increase in anticipatory lick rates for both CS+ and CS-on Days 1 and 2, which remained elevated throughout learning (**Fig. 5h**). Notably, the anticipatory lick rate for CS-was significantly higher than in control mice (**Fig. S4b**). Furthermore, similar to S1 inactivation using muscimol, silencing either IT or ET neurons had no effect on the behavioral performance of expert mice (**Fig. S5**). These results suggest distinct functional roles for IT and ET neurons in shaping stimulus-reward associative learning, where IT neurons are critical for the association to be formed, and ET neurons are critical for CS discrimination to be refined.

### A reinforcement learning model reflecting L5 subnetworks captures learning dynamics

The differences in response properties in IT and ET neurons and their impact on learning imply distinct computational roles in learning stimulus-reward associations. To explore this further, we employed mathematical modeling to decipher their specific contributions. The Rescorla-Wagner (RW) model is a simple yet widely used reinforcement learning framework for understanding classical Pavlovian conditioning^32^. The basic principle of the RW model is to update the value function (i.e., association strength) for each conditioned stimulus by calculating the reward prediction error (RPE) through learning. We implemented an RW-type model within a neural network framework, in which the neural network was trained to predict the value of each stimulus type (CS+ vs. CS-) using an RW update rule, here referred to as ‘value-encoding network’ (**Fig. 6a**). Next, we extended this model to incorporate three key features observed in our experimental results. First, IT neurons, which exhibited stable representations of stimuli before learning (**Figs. 2, 3**), were modeled as a neural network that was pre-trained in an unsupervised learning task to reconstruct (see Methods) and remained unchanged during stimulus-reward training. After unsupervised training, the IT neural network developed representations that help differentiate between stimuli, thus helping the value-encoding network in predicting the reward values (**Fig. 6a, left**). Second, based on our experimental results showing that reward-predicting responses to stimuli evolve in ET neurons over learning (**Fig. 3**), we modeled ET neurons as relaying reward prediction signals generated by the value-encoding network to an RPE calculation circuit (**Fig. 6a, right**). These signals are then compared with actual rewards, generating RPEs that update the value-encoding network during reward-based training. Lastly, since expert mice were able to perform the task without S1 (**Fig. 1g**), the learned associations (i.e., stimulus values) must therefore be stored in brain regions outside of S1. These regions should therefore also receive sensory input and are capable of executing learned responses independently of S1 during expert-level performance. To model this, the value-encoding network includes two input channels: ‘IT’ and ‘non-S1’ (**Fig. 6a, left**). Both channels independently transmit sensory information to predict stimulus values. This component of the model predicts that the IT channel is responsible for conveying richer stimulus representations coming from the IT network, while the non-S1 channel only conveys coarse stimulus representations (see Methods).

**Figure 6.**
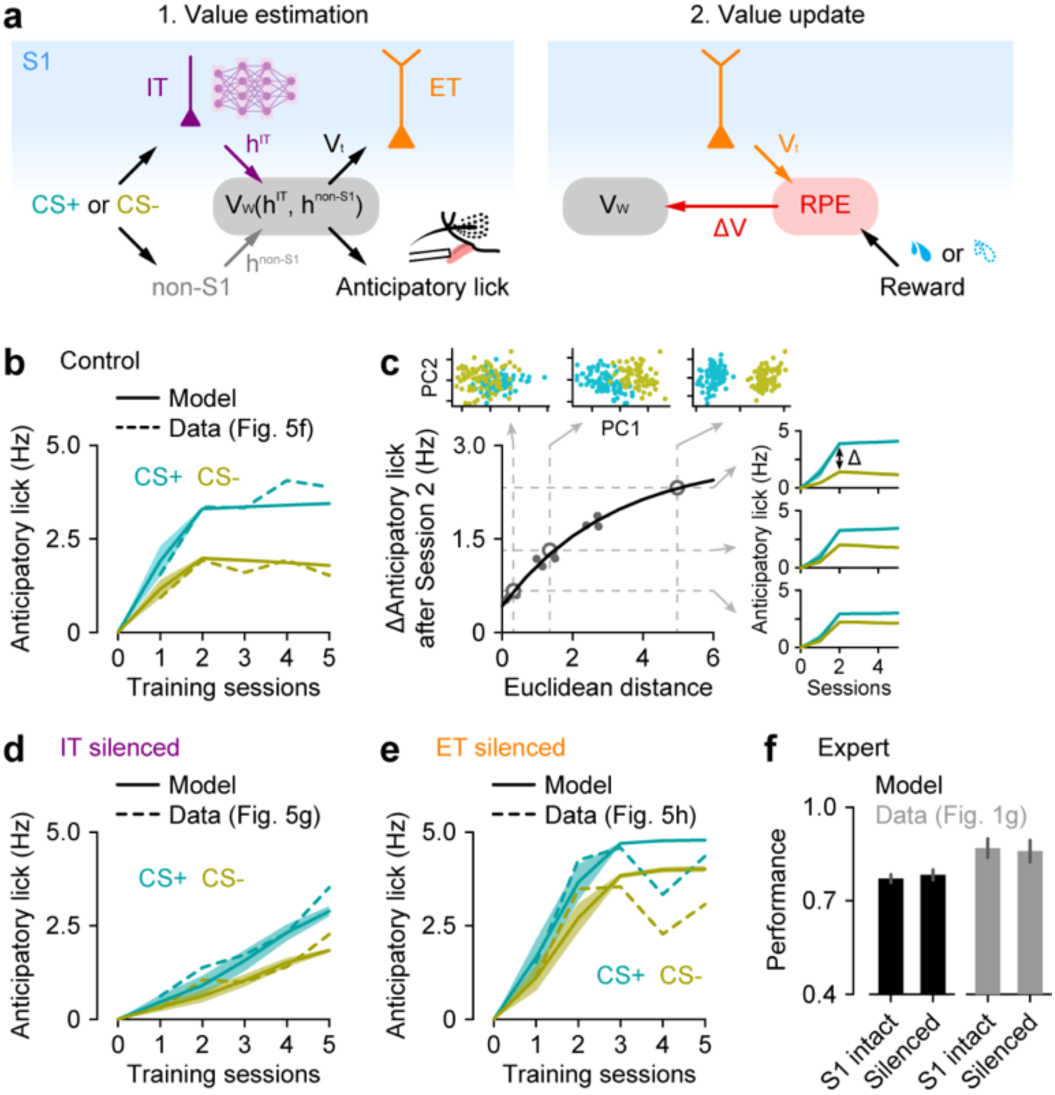
A reinforcement learning model of L5 subnetworks reproduces learning dynamics across conditions. (a) Schematics of the Rescorla-Wagner (RW)-type model, which implements two steps: value estimation and value update. Left, a pre-trained IT network (purple) processes the stimulus, providing a stimulus representation (*h^IT^*), with which the value-encoding network (*V_W_*) estimates the value of the stimulus (*V_t_*). Additionally, the value-encoding network also receives a stimulus representation, *h^non-IT^*, independently of S1. The predicted value is also used to model the anticipatory lick rate. Right, ET neurons convey the predicted value, which is then compared with the actual reward to calculate the reward prediction error (RPE). Finally, the RPE is sent back to the value-encoding network to update the prediction function. (b) Association strengths (solid lines) from the model for CS+ and CS-stimuli through training in the control condition, plotted together with anticipatory lick rates (dashed lines) from the experimental data. Data are presented as mean ± SEM. (c) Relationship between learning dynamics and overlap between CS+ and CS-stimuli in the IT network representational space (black dots), with an exponential fit (black line). Learning dynamics is represented by the difference in association strengths (i.e., anticipatory licks) between distinct CS+ and CS-pairs after Session 2. Representational overlap is measured as the Euclidean distance between CS+ and CS-in the IT network space. Inset, the first two principal components (PCs) of IT representations for three example CS+ and CS-pairs, marked as open circles in the main panel, and their corresponding association strengths across training. (d) Same as **b** but for the condition where IT neurons were silenced. (e) Same as **b** but for the condition where ET neurons were silenced. (f) Discrimination performance in experts before and after silencing of S1, from the model (black) and the experimental data (gray). Data are presented as mean ± SEM.

The model was trained to associate two distinct classes of artificial stimuli, representing CS+ and CS-, with the correct reward outcome. The predicted anticipatory lick rate was derived by linearly scaling the association strength, or value, of each stimulus. Comparison with experimental data showed that the model accurately reproduced the learning dynamics across conditions. Specifically, in the control condition, the model closely tracks the tendency in the experimental data for the initial rise in lick rates for both stimuli, followed by a divergence where CS+ licks continued to increase, and CS-licks declined (**Fig. 6b**). Analysis of the IT neural network in the model revealed that this early general increase could be attributed to the proximity of CS+ and CS-stimuli in the encoding space (**Fig. 6c**), suggesting that overlapping stimulus representations in IT neurons contribute to the general increase observed in the experiments (**Fig. 5f**). The model also captured the distinct learning dynamics observed when IT or ET neurons were blocked. Because in the model IT neurons are responsible for rich sensory encoding, blocking them resulted in a general slower learning for both stimuli (**Fig. 6d**). On the other hand, because ET neurons mediate the transmission of value for RPE computations blocking highlights the fast sensory learning mediated by IT neurons, but inability to correctly assign values to specific stimuli (**Fig. 6e**). Therefore, our model shows that comparing ET neuronal output against the actual reward outcome to generate a RPE is consistent with our data (**Fig. 5h**). Furthermore, it indicates the potential involvement of an RPE calculation circuit downstream of ET neurons. Finally, as expected, our model exhibits learning transfer, so that as learning progresses, the non-S1 input channel in the value-encoding network increasingly builds stimulus-reward associations, leading to a reduced reliance on the IT input channel. This aligns with our experimental observation that, at the expert level, neither IT nor ET neurons in S1 are required for task execution (**Fig. 6f**).

In conclusion, the model effectively captured the distinct contributions of IT and ET neurons to stimulus-reward learning and accurately mirrored experimental findings. These results highlight the complementary computational roles of IT and ET neurons, which fit within a classical associative learning model, where IT neurons provide representations of sensory stimuli to estimate stimulus value, and ET neurons relay this value to the prediction error regions for updating the value function.

## DISCUSSION

Recent studies have begun to reveal distinct roles for L5 IT and ET neurons in cortical sensory areas in decision-making, perception, and cognitive processes^4,5,16,17,21^. Our study extends the understanding of these roles by demonstrating the unique contributions of each L5 subtype to stimulus-reward associative learning. Specifically, we found that IT neurons have stable and robust responses to the stimuli from the beginning which are maintained throughout the learning process, regardless of the associated stimulus value. This indicates their critical role in reliably encoding sensory information. In contrast, ET neurons showed dynamic changes, with their activity patterns evolving as learning progressed. Their responses to the stimuli became more pronounced, coinciding with the emergence of anticipatory licking, suggesting that ET neurons encode the value and behavioral relevance of a stimulus, rather than merely responding to sensory input.

Our chemogenetic inactivation experiments corroborated these distinctions: silencing IT neurons severely impaired the development of anticipatory licking for both CS+ and CS-, underscoring their essential involvement in the association of cue with reward. On the other hand, silencing ET neurons did not affect the overall increase in anticipatory licking but specifically prevented the suppression of licking to CS-, demonstrating their role in refining and differentiating learned responses.

Moreover, these findings align seamlessly with the classical reinforcement learning framework. Our modeling results suggest that IT neurons encode sensory stimuli based on a pre-acquired world model, facilitating value estimation that is crucial for learning. In contrast, ET neurons are engaged in transmitting this estimated value to compute prediction errors, a process fundamental to refining associations. Together, we conclude that L5 IT and ET neuronal subpopulations in the sensory cortex contribute in distinct yet complementary ways to stimulus-reward associative learning.

### Involvement of S1 in stimulus-reward learning

Pharmacological inactivation of S1 severely impaired learning (**Fig. 1f**), while it did not disrupt the performance in expert mice (**Fig. 1g**). This observation aligns with previous studies, indicating that sensory cortex is critical during learning but not necessarily required for the performance of well-established, habitual responses, particularly for simple sensory tasks^5,29,33^. Importantly, our results imply that learned stimulus-reward associations are maintained outside of S1.

The striatum, a central region involved in reward-based learning^34^, emerges as a likely candidate for the retention of learned associations, as lesions in this area have been shown to abolish conditioned responses even in fully trained animals^29^. Both IT and ET neurons in the sensory cortex project to the striatum. Previous studies have shown that stimulus-reward learning induces synaptic strengthening in cortico-striatal projections^3^ and blocking these pathways disrupts learning^5^, but until this study the specific roles of IT and ET neurons in this process had not been investigated. Given the results of this study, including the fact that IT and ET neuronal output are quite dissimilar, we hypothesize that they each exert different effects on striatal activity during learning, particularly in the context of value estimation and updating of learned associations.

### Contribution of IT neuronal activity to stimulus-reward learning

What are the implications of the clearly distinct response patterns and changes seen in IT vs. ET neurons? A large proportion of the IT neurons responded to the stimuli and were able to distinguish the stimulus types with high accuracy, and their response patterns remained largely unchanged during learning. Extrapolating from these observations, we hypothesize that the IT population forms a pre-trained network that encodes various features with high precision. Nevertheless, it was also notable that the majority of stimulus-responding IT neurons responded to both CS+ and CS-(**Fig. 3c**). Our model suggests that this overlap in representations contribute to the generalized stimulus-reward associations that mice initially acquire during learning (**Fig. 6c**). The representational overlap between similar stimuli could promote generalized learning. In some cases of learning, it may reflect a biological strategy to quickly establish a generalized association between stimulus and outcome per se, and then to take a longer time to establish stimulus-specific differences that more closely predict outcome.

### Contribution of ET neuronal activity to stimulus-reward learning

In contrast to IT neurons, ET neurons showed strong responses to the reward itself or to the stimuli that predict reward, with their response patterns dynamically changing as learning progressed. A recent report has highlighted the reward responsiveness of the apical dendrites of L5 neurons in the sensory cortex^35^, and our ET neuron results are consistent with this. Notably, during learning, ET neurons gradually developed reward prediction activity in response to both CS+ and CS-, coinciding with an increase in anticipatory licks. This raises the question of whether ET neuronal activity represents the motor responses accompanying licking. Further analysis of our calcium activity data in ET neurons separating delayed from early lick trials, showed that this activity is at least partially independent of the motor response (**Fig. S2**). This is consistent with a previous study showing that dendrites of L5 neurons do not respond to spontaneous licks unrelated to rewards^35^, supporting the idea that the ET neurons convey reward prediction signals.

Reward prediction signals are fundamental to the process of reinforcement learning, particularly in refining stimulus-specific responses in associative learning. According to classical theoretical models of reinforcement learning^1,36,37^, prediction errors are computed by comparing reward prediction signals with the actual reward availability, reinforcing responses to CS+ and suppressing responses to CS-. The idea that the output of ET neurons is used to calculate the reward prediction error necessary for refining the learned responses is consistent with the finding that the mice with silenced ET neurons failed to learn to discriminate between CS+ and CS-(**Fig. 6a, e**). However, since the initial association was unaffected by the inactivation of ET neurons, it suggests that their output is not critical for generating positive prediction errors, i.e., positive reinforcement. Instead, they are more likely to be involved in the generation of negative prediction errors, thereby suppressing incorrect responses, such as licking to CS-. Identifying the downstream circuits that compute prediction errors using the output of ET neurons would be an important area for future research. One potential target is the subcortical dopaminergic circuits^38^. Recent reports indicate that transient drops in subcortical dopamine signals are responsible for negative prediction errors and contribute to the attenuation of CS-responses in discrimination learning^39^. The reward prediction signals generated using the output of ET neurons may contribute to these dopamine dips.

How reward prediction signals are generated in ET neurons remains to be investigated in future research. The apical dendrites of L5 neurons, located in cortical layer 1 (L1), are hypothesized to function as critical sites for cortical association mechanisms^40–44^. L1 receives non-sensory top-down inputs from higher cortical areas which may contribute to the formation of reward prediction signals. The orbitofrontal cortex (OFC) is an important structure in value encoding during stimulus-reward associations^45–47^. A recent study by Liu et al. demonstrated that top-down inputs from the OFC are involved in the formation of reward prediction activity within the sensory cortex by modulating the activity of dendrite-targeting inhibitory neurons^48^. Inhibition of top-down inputs from the OFC during learning prevented mice from suppressing incorrect lick responses to reward-irrelevant stimuli, paralleling our results from ET neuronal suppression. Thus, the pathway from the OFC to L5 ET neurons in the sensory cortex may convey reward prediction signals to downstream areas, facilitating the generation of negative prediction errors, essential for refining behavioral responses.

In conclusion, we show that distinct subtypes of L5 neurons in sensory cortex are involved in learning, albeit in different aspects of learning. Overall, IT neurons showed stable responses conveying stimulus information necessary for forming general associations between stimuli and reward. In contrast, ET neuronal responses evolved over learning, conveying reward expectation signals, which were necessary for learning to discriminate between stimuli. Their unique activity patterns highlight parallel cortical computations during learning and demonstrate distinct yet complementary contributions of IT and ET neurons to associative learning^12^.

## ACKNOWLEDGMENTS

We thank Andrada-Maria Marica for helpful discussions for modeling; Mario Carta and Richard Naud for their comments on an earlier version of the manuscript. This study was supported by CNRS (to N.T.), the University of Bordeaux (2020 IdEx Junior Chair to N.T.), Conseil régional Nouvelle-Aquitaine (Bordeaux Neurocampus Junior Chair to N.T.), the ATIP-Avenir program (to N.T.), Fondation Schlumberger pour l’Education et la Recherche (FSER202401018842 to N.T.), Fondation pour la Recherche Médicale (EQU202403018077 to N.T.), Brain Science Foundation (to N.T.), Research Foundation for Opto-Science and Technology (to N.T.), the EINSTEIN Foundation Berlin (PhD fellowship to S.M.; A-2021-644 to A.G.; EZ-2014-226 to D.S.), the Deutsche Forschungsgemeinschaft (EXC-2049 – 390688087, LA 3442/3-1, LA 3442/6-1, 327654276/SFB1315 to M.E.L.), the European Union Horizon 2020 Research and Innovation Programme (SGA1-3: 72070/HBP, 785907/HBP, 945539/HBP, 670118/ERC ActiveCortex, 101055340/ERC Cortical Coupling to M.E.L.), the Wellcome Trust (S122871-115 Transition Fellowship to M.G.), the BBSRC (BB/X013340/1 to R.P.C.), EPSRC (EP/X029336/1 to R.P.C.), and the ERC-UKRI Frontier Research Guarantee Starting Grant (EP/Y027841/1 to R.P.C.). We thank the colleagues of the Research Workshop at the Charité - Universitätsmedizin Berlin for developing and manufacturing the experimental devices.

## AUTHOR CONTRIBUTIONS

S.M., M.E.L. and N.T. conceived the project. S.M. and N.T. designed the experiments. S.M. performed experiments. S.M. and C.M. performed the data analysis. M.G. and R.P.C. performed computational modeling. S.M. and N.T. wrote the paper with comments from the other authors.

---

**Figure S1.**
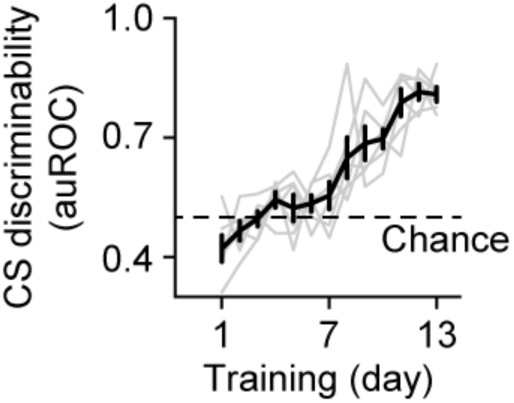
Mice can learn the reversed paradigm. Evolution of behavioral discrimination between CS with reversed CS configuration (CS+ is 5 Hz and CS-is 10 Hz) over 13 days of training (*n* = 6 mice; *p* = 7.1 × 10^-17^, *F* = 20.45; one-way repeated-measure ANOVA). Gray lines, individual mice. Data are presented as mean ± SEM.

**Figure S2.**
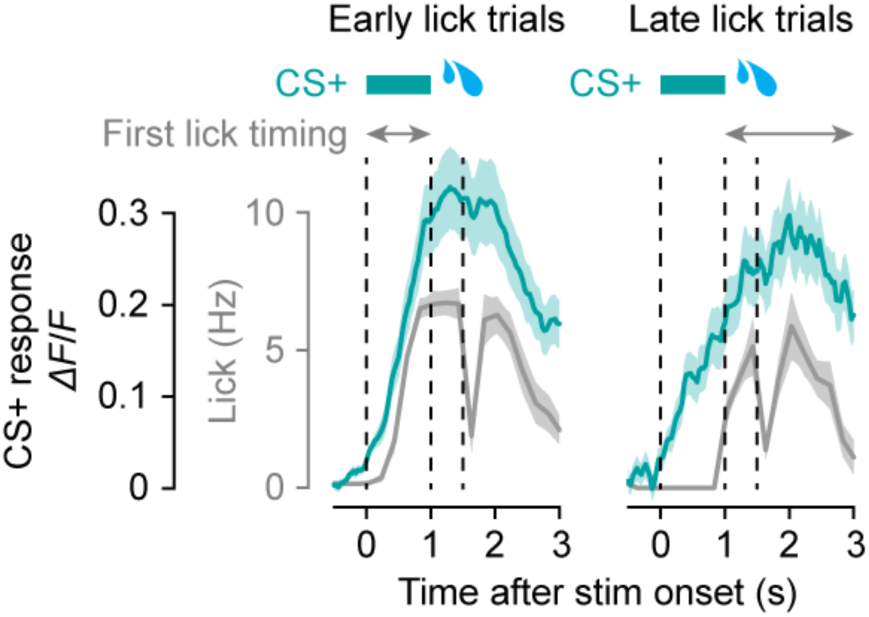
ET neuronal signals are not a simple reflection of lick-related motor activity. Left, calcium responses in CS+ trials on Day 5, where mice initiated licking during the stimulus presentation time window, averaged across ET neurons responding to CS+ (*n* = 71 neurons from 6 mice; 75–92 early lick trials per mouse). Right, calcium responses for CS+ trials where mice licked after the stimulus presentation time window (*n* = 71 neurons from 6 mice; 8–25 late lick trials per mouse). Note that there was calcium activity in the stimulus window despite the lack of licking. Blue trace, ET neuronal calcium responses; gray trace, average lick rates. Data are presented as mean ± SEM.

**Figure S3.**
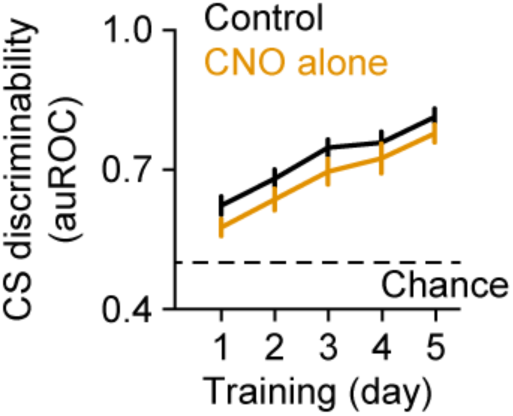
CNO alone does not disrupt the learning progression. Behavioral performance of control mice with intact S1 (*n* = 20 mice) and wild-type mice injected with CNO (*n* = 6 mice) (*p* = 0.14, *F* = 2.34; two-way repeated-measure ANOVA with post hoc Tukey-Kramer test). Data are presented as mean ± SEM.

**Figure S4.**
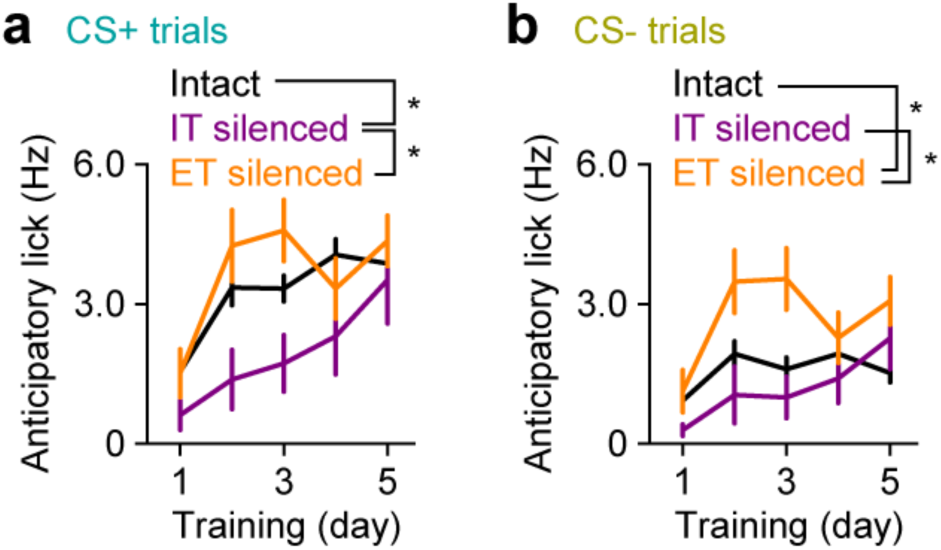
IT and ET neuronal inactivation affects the evolution of anticipatory licking through learning. (a) Anticipatory lick rates for CS+ trials for mice with intact S1 (*n* = 20 mice), mice with silenced IT neurons, (*n* = 6 mice) and mice with silenced ET neurons (*n* = 6 mice; *p* = 0.031, *F* = 3.92; two-way repeated measure ANOVA with post hoc Tukey-Kramer test). \**p* < 0.05. Data are presented as mean ± SEM. (b) Anticipatory lick rates for CS-trials for mice with intact S1 (*n* = 20 mice), mice with silenced IT neurons, (*n* = 6 mice) and mice with silenced ET neurons (*n* = 6 mice; *p* = 0.017, *F* = 4.71; two-way repeated measure ANOVA with post hoc Tukey-Kramer test). \**p* < 0.05. Data are presented as mean ± SEM.

**Figure S5.**
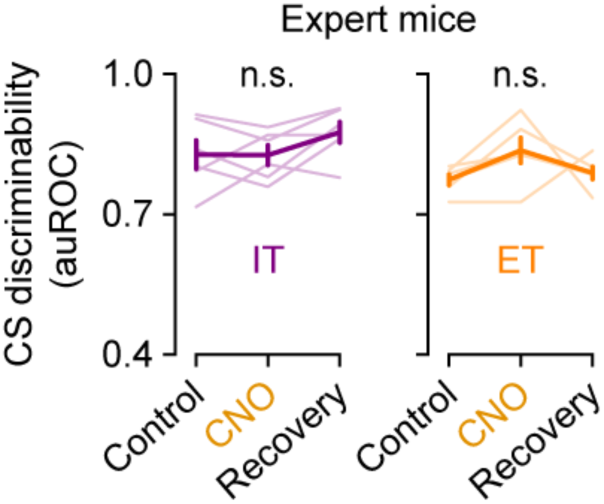
IT and ET neuronal inactivation does not disrupt behavior after learning. Behavioral performance of expert mice with silenced IT neurons (*n* = 6 mice; *p* = 0.077, *F* = 3.36; left) and silenced ET neurons (*n* = 6 mice; *p* = 0.13, *F* = 2.56; one-way repeated-measure ANOVA; right). Gray lines, individual mice. Data are presented as mean ± SEM.

## METHODS

### Animals

Adult C57BL/6J wild-type mice (male and female) or Sim1-Cre (KJ18) (MMRRC, no. 031742-UCD) and Tlx3-Cre (PL56) (MMRRC, no. 041158-UCD) transgenic mice (male and female) mice (>P60) were used. Mice were housed in groups of 2–4 mice per cage in a 12:12 reversed day-night cycle. All experiments were conducted following the guideline given by Landesamt für Gesundheit und Soziales Berlin (LAGeSo) and were approved by this authority.

### Surgical procedures

During the surgery, mice were anesthetized with isoflurane (1.5–2.0% in O_2_) or ketamine (100 mg/kg)/xylazine (10 mg/kg) and kept on a thermal blanket. The skin was removed and the skull was carefully cleaned and scraped with a surgical scalpel. A light-weight head-post was fixed on the skull over the right hemisphere with light-curing adhesives and dental cement. Intrinsic imaging was performed over the skull under light isoflurane anesthesia (1.5-0.8% in O_2_) to locate the C2 barrel column for viral injections and pharmacology.

For chemogenetic inhibition, AAV2/1-hSyn-DIO-hM4D(Gi)-mCherry (Addgene, Product #44362) was injected through a glass pipette (tip diameter, 5–10 µm) into the contralateral (left) S1 (100 nL at 700 μm depth under the pia). The experiments started 3 weeks after viral injection. For *in vivo* two-photon calcium imaging, AAV2/1-syn-FLEX-jGCaMP8m-WPRE (Addgene, Product #162378) was injected through a glass pipette (tip diameter, 5–10 µm) into the contralateral (left) C2 barrel column (100 nl at 700 μm depth under the pia). A 3-mm diameter craniotomy was made over the injected area and the skull and dura were carefully removed. The craniotomy was sealed with a triple-layered glass coverslip (3 mm diameter for the two bottom layers and 4 mm diameter for the top layer) with cyanoacrylate glue and dental cement. Experiments started 3 weeks after viral injection.

### Behavior

Mice were kept on a reversed light/dark cycle. Habituation of the mice to head restraint began a week after the head-post surgery. Head-restrained time on the first day was 5 min and then gradually increased each day until the mice sat calmly for an hour. Mice were water restricted during habituation and subsequent periods of behavioral training.

Behavioral events (e.g., licking, whisker deflection, reward delivery) were monitored and controlled by a custom-written program running on a microcontroller board (Arduino). The C2 whisker was deflected by displacing a light metal tube (∼3 mg) slid over the whisker using a magnetic coil placed underneath the animal. Local magnetic force was generated by loading the coil with a sinusoidal current. Mice were exposed to one of two different frequencies of whisker deflection, 10 Hz and 5 Hz, with a duration of 1s. The 10 Hz whisker deflection (CS+) was paired with a water reward (∼5 µl), delivered through a lick port 0.5 seconds after the stimulus offset, while the 5 Hz whisker deflection (CS-) was not followed by any reward. Licking was detected by a piezo-electric device attached to the licking spout. The mice were allowed to lick at any time, and no punishments were given for any premature licking event. Each mice received 100 trials of each stimulus type (200 trials in total, each trial having an inter-trial interval of 6–8 s), each session repeated for 5 days. The probability of receiving CS+ or CS-was 50% during each session.

For *in vivo* calcium imaging, there was a pre-training and post-training imaging session one day prior as well as one day after training respectively. In these sessions mice were exposed to the stimuli only (without reward) and reward alone (without stimuli) to assess the stimulus response and the reward response, respectively, of the neurons before and after training.

### Behavioral performance analysis

To assess the performance of the task, we used the anticipatory licking occurring from the stimulus onset to the time of the reward (1.5 s) to calculate the area under a receiver operating curve (auROC). auROC ranges from 0 to 1 where auROC > 0.5 indicates higher amount of anticipatory licks in the CS+ trials and auROC < 0.5 indicates higher amount of anticipatory licks in the CS-trials.

### *In vivo* pharmacology

Before every training session, the mice were lightly anesthetized with isoflurane (1.5–2.0% in O_2_). A very small craniotomy was made above the C2 barrel column and muscimol (5 mM, Tocris) was injected at two depths (at depths of 700 µm and 350 µm, 100 nl each). The craniotomy was sealed with a silicone sealant (Kwik-Cast, World Precision Instruments). The mice were put back in their homecage to recover for 10 min before commencing the behavior. After the last training session of the pharmacology experiment, fluorescent muscimol (BODIPY TMR-X conjugate, Thermo Fisher Scientific) was injected into the same injection site (at depths of 700 µm and 350 µm, 100 nl each), and mice were perfused 30 min after the injection.

### *In vivo* chemogenetics

The experiments started 3 weeks after viral injection of hM4Di. Before each session, mice were injected with CNO (5 mg/kg intraperitoneally, Tocris) under brief anesthesia with isoflurane and kept for 30 min in a home cage prior to behavioral testing.

### Histology

Mice were anesthetized using isoflurane (1.5–2% in O_2_) and euthanized by an intraperitoneal injection of urethane (1.5 g/kg). Mice were perfused transcardially with 0.1 M PBS, followed by 4% paraformaldehyde (PFA) in PBS. After perfusion, brains were removed from the skull and postfixed in PFA overnight. The next day, brains were washed in PBS, transferred into a 30% sucrose solution in PBS, and left for 24–48 h for cryoprotection. For cryosectioning, brains were embedded in optimal cutting temperature compound. Coronal brain sections (70 μm) were washed twice in PBS for a minimum of 10 min each at room temperature before staining the nuclei were stained using DAPI (NucBlu Fixed Cell ReadyProbe Reagent, ThermoFisher). After washing, sections were mounted on slides and coverslipped with Fluoromount-G mounting medium. Images were obtained using a fluorescent microscope (DMI 4000B, Leica Microsystems).

### Two-photon calcium imaging

Imaging from behaving mice was performed with a resonant-scanning two-photon microscope (Thorlabs) equipped with GaAsP photomultiplier tubes (Hamamatsu). jGCaMP8m was excited at 940 nm with a Ti:Sapphire laser (Mai Tai eHP DeepSee, Spectra-Physics) and imaged through a 16×, 0.8 NA water-immersion objective (Nikon). Full-frame images (512 × 512 pixels; pixel size, 0.35 × 0.35 µm^2^) were acquired from the apical trunks of ET or IT neurons expressing jGCaMP8m at a depth from the pia of ∼200 µm at 30 Hz using ScanImage 4.1 software (Vidrio Technologies).

### Imaging data analysis

Motion correction of raw files and regions of interest (ROI) selection was performed using Suite2p^49^. Fluorescence change (Δ*F*/*F*_0_) was calculated for each trial where *F*_0_ was the average fluorescence during trial baseline, i.e., during 1 s prior to stimulus onset. ROIs were considered stimulus responsive if the time-averaged response during 1.5 s after stimulus onset significantly increased than the baseline activity during 1 s prior to stimulus onset (paired two-sided Wilcoxon signed rank test, alpha = 0.01), and if their normalized response exceeded 0.05 Δ*F*/*F*_0_. For reward alone trials, we compared the baseline activity to the mean activity during 1.5 s after reward delivery.

We trained a linear Support Vector Machine (SVM) classifier to assess if we could predict trial type (CS+ vs. CS-trials) from the neuronal data. We trained the SVM on 80% of the trials and tested it on the residual 20% of the trials. Trials were binned at 500 ms and the SVM performance was evaluated for each bin. This process of random resampling of train and test data was repeated 1000 times for each bin and the performance of the decoder was collected after each iteration.

Stability of individual neuronal response was measured by performing a Pearson correlation coefficient for each neuron across the 5 days. To assess both the stimulus response stability and reward response stability, trial average responses in the time window of 3 s from stimulus onset to 1.5 s after reward delivery.

### Computational model

We designed a Rescorla Wagner-type model^50^, which learned the value (association strength) of each stimulus (CS+ vs. CS-) over multiple trials.

We modeled the encoding of different stimuli values, i.e., association strengths, using a simple feedforward ‘value-encoding neural network’ denoted as *V_w_* with synaptic weights, *w*. This network consisted of a single linear hidden layer with 116 units where the network inputs were combined into a unified representation, which was then fed to an output layer with sigmoid activation. The value-encoding network, *V_w_*, consisted of two distinct input channels: an S1 IT input channel, providing the IT neuronal representation of the current stimulus (i.e., CS+ or CS-stimuli) and a non-S1 input channel, providing a ‘raw’ representation of the current stimulus independent of S1. At each trial *t*, the value-encoding network prediction for the current stimulus, *x*_t_, is defined as,

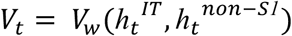

where *h*_t_^IT^ denotes the IT representation of the current stimulus, *x*_t_, while *h*_t_^non-SI^ denotes the non-S1 representation of the same stimulus, *x*_t_. The IT representation, *h*_t_^IT^, was obtained from a separate neural network, IT network, which was pre-trained with unsupervised learning (see below). Conversely, we assumed the non-S1 representation, *h*_t_^non-SI^, to be the ‘raw’ representation of the stimulus, *x*_t_(i.e., *h*_t_^non-SI^ = *x*_t_). Note, the same non-S1 representation *h*_t_^non-SI^, was also used by a separate value-encoding sub-network to support S1-independent expert performance, as described below.

The value-encoding network adapted its synaptic weights, *w*, to predict the correct values, V, for CS+ and CS-stimuli based on the experienced reward outcomes. Following RW models, the value-encoding network synaptic weights were updated at each trial *t* as following,

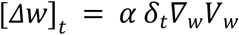

where δ_t_ denotes the reward prediction error (RPE) at trial *t*,

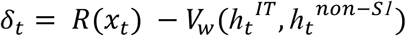

Here, *R* denotes the reward for the current stimulus, *x*_t_. Note that we assumed the reward to be 1 for CS+ stimuli, while to be 0 for CS-stimuli, i.e., *R*(*x*_t_ = *CS+*) = *1* and *R*(*x*_t_ = *CS*−) = *0*.

In summary, at each trial *t*, the value-encoding network, V_w_, took the IT stimulus representation, *h*_t_^IT^, as well as the non-S1 stimulus representation, *h*_t_^non-SI^, as inputs and was trained to predict the correct value, V_w_, for the current stimulus, *x*_t_. Finally, the value-encoding network included a separate sub-network, which could predict the value of each stimulus exclusively based on the non-S1 representation, *h*_t_^non-SI^. During learning, this network was trained based on the S1 IT-dependent value predictions, V_w_(*h*_t_^IT^, *h*_t_^non-SI^), using a simple mean squared loss. In this way, at expert performance, the model could solve the task without requiring S1 IT stimuli representations.

### IT network pre-training

The IT neural network consisted of a feedforward auto-encoder architecture with two encoder layers (with 50 units each), a bottleneck layer (with 15 units) and two decoder layers (with 50 units each). This IT network was pre-trained via unsupervised learning, using a standard L2 loss between the original and the network-reconstructed stimuli^51^. The stimuli were generated by sampling from eight distinct 20-dimensional Gaussian distributions with distinct random means and equal isotropic covariances. Each Gaussian distribution represented a different stimulus type, such as a specific whisker stimulation frequency, with each sample representing a noisy representation of the corresponding stimulus type. Note, we used Gaussian distributions to represent different stimulus types rather than fixed values to account for noise in sensory processing. Two of the eight distinct types of stimuli were randomly selected to be used as CS+ and CS-stimuli in the stimulus-reward association task. Once the IT network was pre-trained, its synaptic weights were fixed and did not change during the association task. The IT-dependent representation, *h*_t_^IT^, for the current stimulus, *x*_t_, was derive as the output of the second encoder layer.

### IT and ET silencing

We modeled IT chemogenetic silencing by replacing the IT-dependent input, *h*_t_^IT^, to the value-encoding network with zero-mean isotropic Gaussian noise. As a result, the value-encoding network could no longer exploit the IT-dependent stimuli representations to estimate the value of each stimulus. Similarly, we reproduced ET chemogenetic silencing by replacing the value predictions relayed by ET neurons with isotropic Gaussian noise. As a result, RPEs could no longer be estimated correctly.

### IT representational overlap and learning performance

For the IT representational overlap analysis, we used the eight Gaussian distributions of stimuli that pre-trained the IT network. Randomly pairing these distributions allowed us to assess how overlap between IT representations of stimulus pairs (CS+ vs. CS-) influences learning in the stimulus-reward association task. We computed the mean Euclidean distance between IT representations, *h*_t_^IT^, across paired stimuli and measured learning performance as the difference in stimulus association strength (i.e., value prediction) between CS+ and CS-after Session 2, where a larger positive difference indicated better learning.

## REFERENCES

1. Watabe-Uchida, M., Eshel, N. & Uchida, N. Neural Circuitry of Reward Prediction Error. Annu Rev Neurosci 40, 373–394 (2017).

2. Moore, S. & Kuchibhotla, K.V. Slow or sudden: Re-interpreting the learning curve for modern systems neuroscience. IBRO Neurosci Rep 13, 9–14 (2022).

3. Xiong, Q., Znamenskiy, P. & Zador, A.M. Selective corticostriatal plasticity during acquisition of an auditory discrimination task. Nature 521, 348–351 (2015).

4. Tang, L. & Higley, M.J. Layer 5 Circuits in V1 Differentially Control Visuomotor Behavior. in Neuron 346–354.e345 (Cell Press, 2020).

5. Ruediger, S. & Scanziani, M. Learning speed and detection sensitivity controlled by distinct cortico-fugal neurons in visual cortex. in eLife 1–24 (eLife Sciences Publications Ltd, 2020).

6. Letzkus, J.J., Wolff, S.B., Meyer, E.M., Tovote, P., Courtin, J., Herry, C. & Luthi, A. A disinhibitory microcircuit for associative fear learning in the auditory cortex. Nature 480, 331–335 (2011).

7. Henschke, J.U., Dylda, E., Katsanevaki, D., Dupuy, N., Currie, S.P., Amvrosiadis, T., Pakan, J.M.P. & Rochefort, N.L. Reward Association Enhances Stimulus-Specific Representations in Primary Visual Cortex. Curr Biol 30, 1866–1880 e1865 (2020).

8. Gilad, A. & Helmchen, F. Spatiotemporal refinement of signal flow through association cortex during learning. Nat Commun 11, 1744 (2020).

9. Esmaeili, V., Oryshchuk, A., Asri, R., Tamura, K., Foustoukos, G., Liu, Y., Guiet, R., Crochet, S. & Petersen, C.C.H. Learning-related congruent and incongruent changes of excitation and inhibition in distinct cortical areas. PLoS Biol 20, e3001667 (2022).

10. Harris, K.D. & Shepherd, G.M. The neocortical circuit: themes and variations. Nat Neurosci 18, 170–181 (2015).

11. Shepherd, G.M.G. & Yamawaki, N. Untangling the cortico-thalamo-cortical loop: cellular pieces of a knotty circuit puzzle. Nat Rev Neurosci 22, 389–406 (2021).

12. Moberg, S. & Takahashi, N. Neocortical layer 5 subclasses: From cellular properties to roles in behavior. Front Synaptic Neurosci 14, 1006773 (2022).

13. Gerfen, C.R., Paletzki, R. & Heintz, N. GENSAT BAC cre-recombinase driver lines to study the functional organization of cerebral cortical and basal ganglia circuits. Neuron 80, 1368–1383 (2013).

14. Tervo, D.G., Hwang, B.Y., Viswanathan, S., Gaj, T., Lavzin, M., Ritola, K.D., Lindo, S., Michael, S., Kuleshova, E., Ojala, D., Huang, C.C., Gerfen, C.R., Schiller, J., Dudman, J.T., Hantman, A.W., Looger, L.L., Schaffer, D.V. & Karpova, A.Y. A Designer AAV Variant Permits Efficient Retrograde Access to Projection Neurons. Neuron 92, 372–382 (2016).

15. Znamenskiy, P. & Zador, A.M. Corticostriatal neurons in auditory cortex drive decisions during auditory discrimination. Nature 497, 482–485 (2013).

16. Takahashi, N., Ebner, C., Sigl-Glockner, J., Moberg, S., Nierwetberg, S. & Larkum, M.E. Active dendritic currents gate descending cortical outputs in perception. Nat Neurosci 23, 1277–1285 (2020).

17. Musall, S., Sun, X.R., Mohan, H., An, X., Gluf, S., Li, S.J., Drewes, R., Cravo, E., Lenzi, I., Yin, C., Kampa, B.M. & Churchland, A.K. Pyramidal cell types drive functionally distinct cortical activity patterns during decision-making. Nat Neurosci 26, 495–505 (2023).

18. Economo, M.N., Viswanathan, S., Tasic, B., Bas, E., Winnubst, J., Menon, V., Graybuck, L.T., Nguyen, T.N., Smith, K.A., Yao, Z., Wang, L., Gerfen, C.R., Chandrashekar, J., Zeng, H., Looger, L.L. & Svoboda, K. Distinct descending motor cortex pathways and their roles in movement. Nature 563, 79–84 (2018).

19. Heindorf, M., Arber, S. & Keller, G.B. Mouse Motor Cortex Coordinates the Behavioral Response to Unpredicted Sensory Feedback. Neuron 99, 1040–1054 e1045 (2018).

20. Li, N., Chen, T.W., Guo, Z.V., Gerfen, C.R. & Svoboda, K. A motor cortex circuit for motor planning and movement. Nature 519, 51–56 (2015).

21. Mohan, H., An, X., Xu, X.H., Kondo, H., Zhao, S., Matho, K.S., Wang, B.S., Musall, S., Mitra, P. & Huang, Z.J. Cortical glutamatergic projection neuron types contribute to distinct functional subnetworks. Nat Neurosci 26, 481–494 (2023).

22. Wang, L.-P., Bodner, M. & Zhou, Y.-D. Distributed Neural Networks of Tactile Working Memory. Journal of Physiology-Paris 107, 452–458 (2013).

23. Romo, R. & Rossi-Pool, R. Turning Touch into Perception. Neuron 105, 16–33 (2020).

24. Staiger, J.F. & Petersen, C.C.H. Neuronal Circuits in Barrel Cortex for Whisker Sensory Perception. Physiol Rev 101, 353–415 (2021).

25. Francioni, V., Padamsey, Z. & Rochefort, N.L. High and asymmetric somato-dendritic coupling of V1 layer 5 neurons independent of visual stimulation and locomotion. Elife 8 (2019).

26. Beaulieu-Laroche, L., Toloza, E.H.S., Brown, N.J. & Harnett, M.T. Widespread and Highly Correlated Somato-dendritic Activity in Cortical Layer 5 Neurons. Neuron 103, 235–241 e234 (2019).

27. Otis, J.M., Namboodiri, V.M., Matan, A.M., Voets, E.S., Mohorn, E.P., Kosyk, O., McHenry, J.A., Robinson, J.E., Resendez, S.L., Rossi, M.A. & Stuber, G.D. Prefrontal cortex output circuits guide reward seeking through divergent cue encoding. Nature 543, 103–107 (2017).

28. Namboodiri, V.M.K., Otis, J.M., van Heeswijk, K., Voets, E.S., Alghorazi, R.A., Rodriguez-Romaguera, J., Mihalas, S. & Stuber, G.D. Single-cell activity tracking reveals that orbitofrontal neurons acquire and maintain a long-term memory to guide behavioral adaptation. Nat Neurosci 22, 1110–1121 (2019).

29. Hong, Y.K., Lacefield, C.O., Rodgers, C.C. & Bruno, R.M. Sensation, movement and learning in the absence of barrel cortex. Nature 561, 542–546 (2018).

30. Zhang, Y., Rozsa, M., Liang, Y., Bushey, D., Wei, Z., Zheng, J., Reep, D., Broussard, G.J., Tsang, A., Tsegaye, G., Narayan, S., Obara, C.J., Lim, J.X., Patel, R., Zhang, R., Ahrens, M.B., Turner, G.C., Wang, S.S., Korff, W.L., Schreiter, E.R., Svoboda, K., Hasseman, J.P., Kolb, I. & Looger, L.L. Fast and sensitive GCaMP calcium indicators for imaging neural populations. Nature 615, 884–891 (2023).

31. Armbruster, B.N., Li, X., Pausch, M.H., Herlitze, S. & Roth, B.L. Evolving the lock to fit the key to create a family of G protein-coupled receptors potently activated by an inert ligand. Proceedings of the National Academy of Sciences 104, 5163–5168 (2007).

32. Rescorla, R.A. & Wagner, A. A theory of Pavlovian conditioning: Variations in the effectiveness of reinforcement and nonreinforcement. (1972).

33. Ceballo, S., Piwkowska, Z., Bourg, J., Daret, A. & Bathellier, B. Targeted Cortical Manipulation of Auditory Perception. Neuron 104, 1168–1179 e1165 (2019).

34. Cox, J. & Witten, I.B. Striatal circuits for reward learning and decision-making. in Nature Reviews Neuroscience 2019 20:*8* 482–494 (Nature Publishing Group, 2019).

35. Lacefield, C.O., Pnevmatikakis, E.A., Paninski, L. & Bruno, R.M. Reinforcement Learning Recruits Somata and Apical Dendrites across Layers of Primary Sensory Cortex. Cell Rep 26, 2000–2008 e2002 (2019).

36. Schultz, W., Dayan, P. & Montague, P.R. A neural substrate of prediction and reward. Science 275, 1593–1599 (1997).

37. Schultz, W. Predictive reward signal of dopamine neurons. J Neurophysiol 80, 1–27 (1998).

38. Schultz, W. & Dickinson, A. Neuronal coding of prediction errors. Annu Rev Neurosci 23, 473–500 (2000).

39. Iino, Y., Sawada, T., Yamaguchi, K., Tajiri, M., Ishii, S., Kasai, H. & Yagishita, S. Dopamine D2 receptors in discrimination learning and spine enlargement. Nature 579, 555–560 (2020).

40. Larkum, M.E., Zhu, J.J. & Sakmann, B. A new cellular mechanism for coupling inputs arriving at different cortical layers. Nature 398, 338–341 (1999).

41. Larkum, M. A cellular mechanism for cortical associations: an organizing principle for the cerebral cortex. Trends Neurosci 36, 141–151 (2013).

42. Doron, G., Shin, J.N., Takahashi, N., Druke, M., Bocklisch, C., Skenderi, S., de Mont, L., Toumazou, M., Ledderose, J., Brecht, M., Naud, R. & Larkum, M.E. Perirhinal input to neocortical layer 1 controls learning. Science 370 (2020).

43. Shin, J.N., Doron, G. & Larkum, M.E. Memories off the top of your head. Science 374, 538–539 (2021).

44. Xu, N.L. How the brain’s primary processing units compute to give rise to intelligence. Nat Rev Neurosci 25, 288 (2024).

45. Schoenbaum, G., Chiba, A.A. & Gallagher, M. Orbitofrontal cortex and basolateral amygdala encode expected outcomes during learning. Nature Neuroscience 1, 155–159 (1998).

46. Tremblay, L. & Schultz, W. Relative reward preference in primate orbitofrontal cortex. Nature 398, 704–708 (1999).

47. Roesch, M.R. & Olson, C.R. Neuronal Activity Related to Reward Value and Motivation in Primate Frontal Cortex. Science 304, 307–310 (2004).

48. Liu, D., Deng, J., Zhang, Z., Zhang, Z.Y., Sun, Y.G., Yang, T. & Yao, H. Orbitofrontal control of visual cortex gain promotes visual associative learning. Nat Commun 11, 2784 (2020).

49. Pachitariu, M., Stringer, C., Dipoppa, M., Schröder, S., Rossi, L.F., Dalgleish, H., Carandini, M. & Harris, K.D. Suite2p: beyond 10,000 neurons with standard two-photon microscopy. *bioRxiv* (2016).

50. Rescorla, R.A. & Wagner, A.R. A theory of Pavlovian conditioning: Variations in the effectiveness of reinforcement and nonreinforcement (Appleton-Century-Crofts, 1972).

51. Bank, D., Koenigstein, N. & Giryes, R. Autoencoders. (ed. L. Rokach, O. Maimon & E. Shmueli) (Springer Cham, 2023).

